# STRPsearch: fast detection of structured tandem repeat proteins

**DOI:** 10.1101/2024.07.10.602726

**Authors:** Soroush Mozaffari, Paula Nazarena Arrías, Damiano Clementel, Damiano Piovesan, Carlo Ferrari, Silvio C. E. Tosatto, Alexander Miguel Monzon

## Abstract

**Motivation:** State-of-the-art prediction methods are generating millions of publicly available protein structures. Structured Tandem Repeats Proteins (STRPs) constitute a subclass of tandem repeats characterized by repetitive structural motifs. STRPs exhibit distinct propensities for secondary structure and form regular tertiary structures, often comprising large molecular assemblies. They can perform important and diverse biological functions due to their highly degenerated sequences, which maintain a similar structure while displaying a variable number of repeat units. This suggests a disconnection between structural size and protein function. However, automatic detection of STRPs remains challenging with current state-of-the-art tools due to their lack of accuracy and long execution times, hindering their application on large datasets. In most cases, manual curation is the most accurate method for detecting and classifying them, making it impossible to inspect millions of structures.

**Results:** We present STRPsearch, a novel computational tool for rapid identification, classification, and mapping of STRPs. Leveraging the manually curated entries in RepeatsDB as the known conformational space of the STRPs, STRPsearch utilizes the latest advancements in structural alignment techniques for a fast and accurate detection of repeated structural motifs in protein structures, followed by an innovative approach to map units and insertions through the generation of TM-score graphs. STRPsearch can serve researchers in structural bioinformatics and protein science as an efficient and practical tool for analysis and detection of STRPs.

**Availability and implementation:** STRPsearch is coded in Python, all the scripts and the associated documentation are available at https://github.com/BioComputingUP/STRPsearch.

**Supplementary information:** Supplementary data are available..

## 1 Introduction

Tandem Repeat Proteins (TRPs) represent a diverse group of proteins featuring repetitive sequence motifs (Kajava and Tosatto 2018). A specialized TRP subset, known as Structured Tandem Repeat Proteins (STRPs) (Monzon *et al*. 2023) is distinguished by the conservation of specific structural motifs rather than mere sequence repetition. In STRPs, the repetitive units are the fundamental structural elements that collectively constitute repeat regions (Di Domenico *et al*. 2013). The proposed classification by Kajava (Kajava 2012) categorizes tandem repeats into five different classes, based on their architectural arrangement and the length of their constituent units. Recent predictions suggest that 50.9% of proteins across all kingdoms of life are composed of at least one TRP region, with a particular enrichment of TRPs in Eukaryotes (Delucchi *et al*. 2020).

TRPs have been shown to be involved in many biological functions and activities. For example, DNA sliding clamps are TRPs which play an essential role in DNA replication (Arrías *et al*. 2023) while leucine-rich repeats (LRRs) make up the extracellular domains of toll-like receptors (TLRs) involved in host immune responses (Leulier and Lemaitre 2008). In recent years, their significance has garnered increasing attention, owing to their implications in health (de Wit *et al*. 2011; Fournier *et al*. 2013) and their application in protein design (Höcker 2014; Brunette *et al*. 2015; Wu *et al*. 2023). On the other hand, with the steady growth of the Protein Data Bank (PDB), storing >217,000 (mar-2024) experimental protein structures, and the huge amount of protein structural models from the recent structure prediction methods such as AlphaFold (Jumper *et al*. 2021) and RoseTTAFold (Baek *et al*. 2021), the scientific community has an unprecedented volume of protein structure data available. This challenges state-of-the-art methods dealing with protein structures.

RepeatsDB (Paladin *et al*. 2020) is the main repository of STRPs annotation and classification. Through manual curation of STRPs on experimental structures, each entry is classified by precisely identifying regions, units and insertions, including the determination of their position and range within the protein structure. As an outcome of this curation effort, RepeatsDB can serve as a ground-truth for the development and fine-tuning of computational tools designed for the study and analysis of STRPs. Different predictors have been developed for the automatic TRP detection from sequence or structure (Delucchi *et al*. 2021; Kamel *et al*. 2021). Particularly, tools such as TAPO (Do Viet, Roche and Kajava 2015) and RepeatsDB-lite (Hirsh *et al*. 2018) aimed to detect repeated regions and units from protein structures. However, their performance represents a bottleneck for large-scale STRP detection given the current structural data available, e.g. in the PDB and the AlphaFoldDB (Varadi *et al*. 2022). Moreover, the manually curated information of STRPs deposited in RepeatsDB is not being exploited by the state-of-the-art methods. Here, we present STRPsearch, a fast method to accurately detect STRPs on protein structures.

## 2 Methods

The algorithm requires one primary input which is a protein structure as the query. It then utilizes two structural repeat libraries to identify repeated structural motifs within the input structure. The libraries are built upon the reviewed entries in the RepeatsDB (dated 2023-05-03), comprising an extensive dataset of PDB chains in which repeat regions and units have been manually curated. This represents a sample of the conformational space and diversity of STRPs. The algorithm utilizes this data through two main libraries, the Representative-Unit-Library (RUL) and the Tri-Unit-Library (TUL), composed of 2460 proteins (different UniProt IDs), 9121 PDB chains and 9502 repeated regions each.

The “representative unit” is a singular repeat unit of a repeat region that exhibits the maximum structural similarity, measured by TM-Score, to other units within that specific region, and is archived in the RUL. Based on the representative unit’s position within the region, two proximate units are chosen that are closest in sequential relation to the representative unit. These three units, when combined, form a tri-unit structure that is then trimmed and stored in the TUL.

In the first step, the algorithm searches for repeated structural motifs in the query structure. This involves structurally aligning the query structure against each tri-unit structure in the TUL. For this purpose Foldseek (van Kempen *et al*. 2023) is used, since it can quickly align the query structure against a database of target structures and provides different probabilistic metrics and similarity scores of the alignments produced. By the end of this stage, STRPsearch determines the likelihood of the presence of different types of repeated structures, setting the stage for the next steps (Figure 1a).

**Figure 1.**
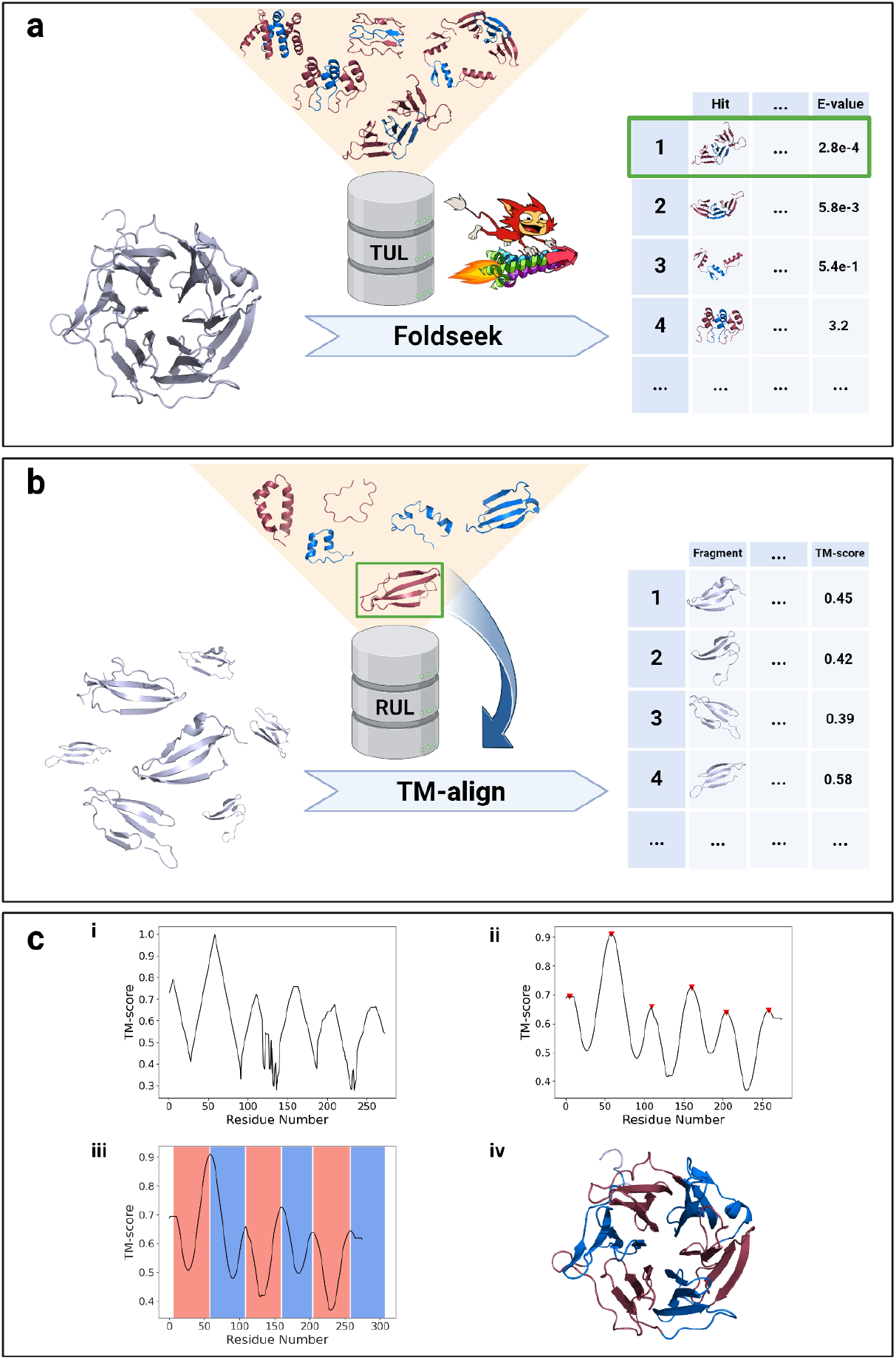
Schematic representation of the STRPsearch software method and its applications. (a) The query structure (chain A of PDB structure 1IUB) is aligned against the Tri-Unit-Library (TUL) using Foldseek; the hits with the lowest E-values will be selected for the subsequent step. (b) The query structure is fragmented into sizes matching the selected hit from the previous step and are aligned against the associated representative unit, retrieved from Representative-Unit-Library (RUL) using pairwise structural alignment software TM-align, yielding pairwise TM-scores for each fragment. (c) [i] TM-scores are plotted against fragment first residues, elucidating relative structural similarity across the whole span of the query structure. [ii] SciPy’s signal toolbox is used to smooth the graph data and detect peaks. [iii] The detected peaks, which signal the potential positions of repeat units, are interpreted by the software to map the repeat region. [iv] The final output visualizes the mapped query structure in PyMOL, with repeat units alternately colored in red and blue.

At the next step, the algorithm utilizes the results from the initial step to select the most probable hit based on the E-value of query-target alignment pairs. In case of co-occurrence of different types of repeats on the same query structure, two or more hits will be selected based on the target classification of probable hits. For every hit, the representative repeat unit associated with its target is retrieved from RUL and is aligned across the full span of the query structure similar to a sliding window approach with a single residue increment at each step, aiming to enhance resolution and accuracy (Figure 1b). To achieve this goal, the query structure is fragmented into pieces matching the length of the target representative unit. These fragments are structurally aligned, in a pairwise manner, to the representative unit using TM-align (Zhang and Skolnick 2005). It quantifies structural similarity between two protein structures using TM-score. The resulting TM-scores are recorded and plotted against the starting residue numbers of the query fragments to create a TM-score profile per residue. These graphs depict the relative structural similarity variations between the query and the representative unit across the whole length of the query structure (Figure 1c).

When repeat regions are present, the alignment of the representative unit with similar repeat units in the region produces periodic peaks in the TM-score profile. To accurately identify these peaks and map the integral components of the repeat region based on their positioning, the TM-score profile, which could be interpreted as a graph, undergoes smoothing adjusted to the length of the representative unit. This optimization enhances the performance of peak detection algorithms. Subsequently, using SciPy’s signal processing toolbox, peaks are identified and the integral components are mapped. If the algorithm identifies a minimum of three adjacent repeat units, it maps a repeat region on the query structure (Figure 1c).

Two key parameters that can be customized via the command line interface are “max_eval” and “min_height”. “max_eval” denotes the upper threshold of E-values for hits identified by Foldseek in the initial phase of the algorithm. A higher “max_eval” leads to an increased incidence of false positive results and conversely, lower values can increase the occurrence of false negatives. “min_height” represents the minimum allowed height of the peaks that are detected by the peak detection method in the TM-score profile, which indicate the potential position of repeat units. “min_height” default values are optimized based on the average structural similarity that is seen among the units of different types of repeats in the RepeatsDB. Other parameters used by default for the peak detection method are optimized based on grid search.

To evaluate the performance of the software, RepeatsDB served as a reference with 2002 positive structures (STRPs) from unique protein sequences containing structural tandem repeats of 6 major types. These types included Alpha-solenoids, Beta-solenoids, Alpha/Beta-solenoids, Beta-propellers, TIM-barrels, and Beta-barrels. A manually curated negative dataset composed of 1737 negative structures (non-STRPs) belonging to unique proteins was also utilized to provide the possibility of false positive and true negative predictions by the software. Subsequently, both the positive and negative datasets were subjected to clustering at 30% sequence identity using BLASTClust (Altschul *et al*. 1990), and one representative entry from each cluster was randomly selected. This process led to a reduction in the size of both the positive and negative datasets. Ultimately, resulting in 1225 positive and 1218 negative structures (see Suppl. Dataset S1 & S2). The evaluation strategy employed a five-fold stratified cross-validation, allowing the algorithm access to 80% of the positive structures as the template conformational space, while the remaining 20%, in conjunction with a non-overlapping 20% segment of the negative dataset, were used for validation.

## 3 Results and Discussion

STRPsearch is developed in Python version 3.8 exploiting various libraries such as Biopython and SciPy. Foldseek, TM-align and PyMOL are external dependencies which are dockerized and integrated in a Conda environment. The source code is provided under the GPL license.

### 3.1 Application on protein structures

To execute the software, there are three options available. The first involves querying a protein structure by providing the input file formatted as PDB/mmCIF, with the option to query a specific chain or all the chains included in the structure. The second option, users can specify the PDB ID allowing the software to automatically download and query a specific chain or all the chains included in the PDB structure. As the third option, the software can directly download and query an AlphaFold model by indicating the UniProt ID.

Upon the identification of STRPs, the output of the algorithm for each identified repeat region includes four components: 1. a JSON formatted file containing the classification of the associated repeat region and the boundaries of units and insertions (if they exist), 2. the trimmed structure of the repeat region in PDB format, 3. a PyMOL (Schrödinger, LLC 2015) session of the repeat region colored based on units and insertions, and 4. a TM-score profile per residue, highlighted with the position and range of the repeat units.

### 3.2 Performance evaluation

The cross-validation demostrates that the predictor consistently performs well, with an average, 80% of total positive structures with STRPs being correctly detected, with a standard deviation of 1.9, and 10% of the negative structures falsely predicted as positive, with standard deviation of 2.57 (see cross-validation results on Suppl. Table S3 and S4). Regarding the capability of the software in identifying the residues involved in structured tandem repeat regions, the algorithm resulted in, on average, an accuracy [(TP + TN) / (TP + FP + TN + FN)] of 0.88 in detecting residues inside the repeat regions, with a precision [TP / (TP + FP)] of 0.91, a recall [or sensitivity; TP / (TP + FN)] of 0.91, and a F1-score [2.TP / (2.TP + FP + FN)] of 0.90, with all metrics showing a standard deviation close to 0.01 (see Suppl. Table S5). Overall, the algorithm demonstrates good performance in discerning nearly the entirety of the repeat regions.

### 3.3 Benchmarking

STRPsearch was assessed in comparison to RepeatsDB-lite and TAPO, two softwares for identification of STRPs that are only available to the public as web services. This comparative assessment focused on the ability of different software to distinguish between STRPs and non-STRPs. The evaluation was performed on a dataset consisting of 244 positive structures, randomly selected but stratified based on classification to ensure an even distribution of different repeat types across all folds, along with 244 negative structures. As shown in Table 1, RepeatsDB-lite and TAPO exhibit relatively high recall rates; however, they lag behind STRPsearch in other metrics. This disparity can be attributed primarily to their suboptimal performance in correctly identifying negative structures resulting in high numbers of false positives. Comparison of execution times indicates notable differences as STRPsearch processed each entry in an average time of 9 seconds, with a standard deviation of 9. Meanwhile, TAPO took an average of 26 seconds per entry, with a standard deviation of 55 and RepeatsDB recorded a relatively much longer processing time of 190 seconds on average, accompanied by a standard deviation of 216 (see Suppl. Figure 3).

**Table 1.**
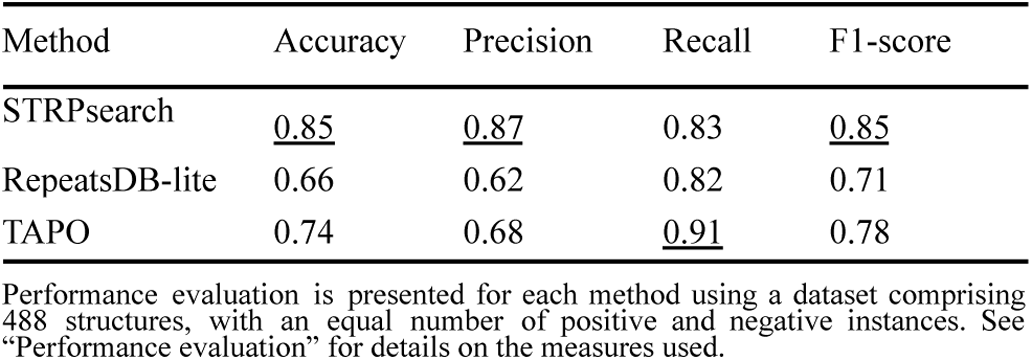
Comparative STRP identification evaluation.

| Method | Accuracy | Precision | Recall | F1-score |
| --- | --- | --- | --- | --- |
| STRPsearch | <u>0.85</u> | <u>0.87</u> | 0.83 | <u>0.85</u> |
| RepeatsDB-lite | 0.66 | 0.62 | 0.82 | 0.71 |
| TAPO | 0.74 | 0.68 | <u>0.91</u> | 0.78 |

Performance evaluation is presented for each method using a dataset comprising 488 structures, with an equal number of positive and negative instances. See "Performance evaluation" for details on the measures used.

### 3.4 Running on PDB and AlphaFoldDB model organism proteomes

Running the software on the PDB, 216,478 protein structures (dated 20/02/2024) resulted in detecting 15,947 structures as STRPs, which corresponds to 4147 unique protein sequences. In another analysis, running on AlphaFoldDB contains structural models of 48 organisms, totaling 564,446 proteins. STRPsearch identified 40,149 proteins as STRPs. While computational runtime is highly correlated with protein structure length, for proteins of a length around 500 residues, the average execution time was 35 seconds with a standard deviation of 20.

## 4 Conclusions/Summary

We presented STRPsearch, a software designed for fast and accurate identification, classification and mapping of structural tandem repeats in protein structures. By exploiting the manually curated entries in RepeatsDB as the ground-truth data and the utilization of the latest computational advancements in the field, the performance metrics of the software outmatches the similar tools by offering heightened reliability, accuracy, and speed. This renders STRPsearch to be a valuable stand-alone tool for identification and further analysis of STRPs that could easily be applied to large protein structure databases.

## Acknowledgements

The authors thank Dr. Andrey Kajava as well as to REFRACT secondees Stefany Neciosup Vera and Hector Hernan Henao Uribe, for their assistance in running TAPO.

