## Supplementary information for "STRPsearch: fast detection of structured tandem repeat proteins"

### Data preparation

To exploit the manually curated dataset of STRPs of RepeatsDB, we employed the most recent update, dated 2023-05-03. Comprising 9834 entries, each entry represents a manually curated repeat region. This dataset is linked to a total of 9448 PDB chains, 5628 PDB IDs, and 2473 UniProt IDs (**Table S1**). Every entry within RepeatsDB is classified into four hierarchical levels following Kajava's classification: Class, Topology, Fold, and Clan, with each subsequent level serving as a subcategory of its predecessor. In our endeavor, we aggregate entries based on the top two levels, namely Class and Topology, to delineate STRP types (**Figure S1**). After selecting STRP types with a sufficient number of entries to evaluate the software's performance, we addressed redundancy by randomly choosing one repeat region for each unique UniProt ID. Subsequently, we clustered at 30% of identity the sequences of the associated PDB chains for the selected samples and randomly chose one sample from each cluster. **Table S2** illustrates the final count of samples for each selected type, resulting in a total of 1225 positive samples.

| RepeatsDB entries | Count |
|---|---|
| Repeat region | 9533 |
| PDB chain | 9140 |
| PDB ID | 5462 |
| UniProt ID | 2473 |

**Table S1.** Number of repeat regions, PDB chains, PDB IDs, and UniProt IDs associated with the entries in the RepeatsDB dataset.

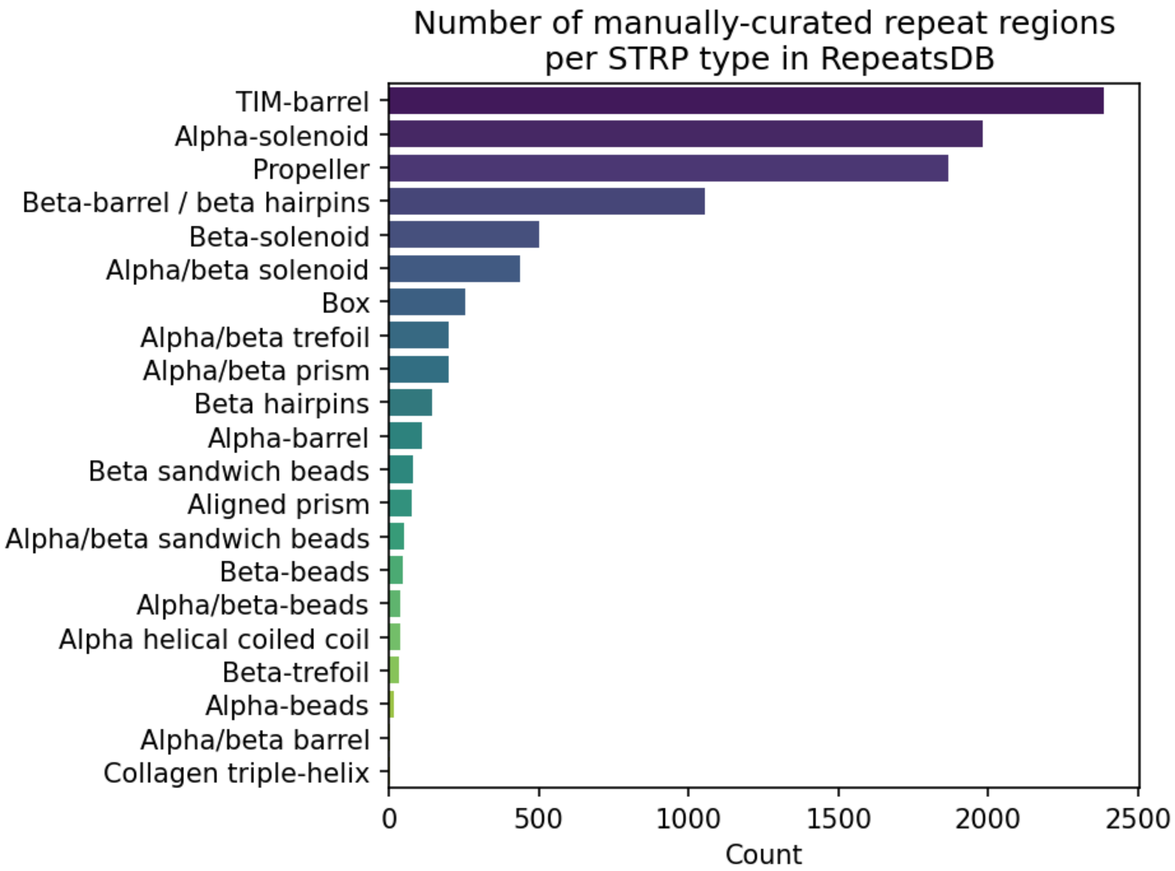


**Figure S1.** Number of entries per each type of STRPs in the RepeatsDB dataset.

| **Type** | **Count** |
|---|---|
| Alpha-solenoid | 401 |
| Propeller | 249 |
| TIM-barrel | 227 |
| Beta-barrel | 133 |
| Beta-solenoid | 120 |
| Alpha/beta solenoid | 95 |

**Table S2**. Number of unique entries per each type of STRP in the dataset used to evaluate the performance of the software.

### Performance results

Following a 5-fold cross-validation, the initial analysis focused on assessing the software's ability to distinguish STRPs from non-STRPs. To achieve this, the confusion matrix was computed for each fold. The raw counts of True Positives (TP), True Negatives (TN), False Positives (FP), and False Negatives (FN) for each fold are shown in **Table S3**. Additionally, performance metrics such as Accuracy, Precision, Recall, and F1-score were calculated to provide a more comprehensive evaluation of the software's performance (**Table S4**). In a separate analysis, we assessed the software's performance in detecting the range of repeat regions by evaluating its ability to identify residues within the curated range of each repeat region. The schematic representation in **Figure S2** shows the rationale behind this approach, and the **Table S5** presents the performance metrics for each fold. **Figure S3** shows the execution time per PDB chain by STRPsearch, RepeatsDB-Lite, and TAPO.

| **Fold** | **TP** | **FP** | **TN** | **FN** |
|---|---|---|---|---|
| 1 | 203 | 31 | 213 | 42 |
| 2 | 195 | 14 | 230 | 50 |
| 3 | 191 | 21 | 223 | 54 |
| 4 | 191 | 23 | 220 | 54 |
| 5 | 191 | 30 | 213 | 54 |

**Table S3**. Number of each component in the confusion matrix computed for each fold in the cross-validation test to evaluate the capability of the software in identifying STRPs from non-STRPs.

| **Fold** | **Accuracy** | **Precision** | **Recall** | **F1-Score** |
|---|---|---|---|---|
| 1 | 0.85 | 0.87 | 0.83 | 0.85 |
| 2 | 0.87 | 0.93 | 0.8 | 0.86 |
| 3 | 0.85 | 0.9 | 0.78 | 0.84 |
| 4 | 0.84 | 0.89 | 0.78 | 0.83 |
| 5 | 0.83 | 0.86 | 0.78 | 0.82 |

**Table S4**. The performance metrics, derived from the confusion matrix calculated for each fold during the cross-validation test, of the software in identifying STRPs from non-STRPs.

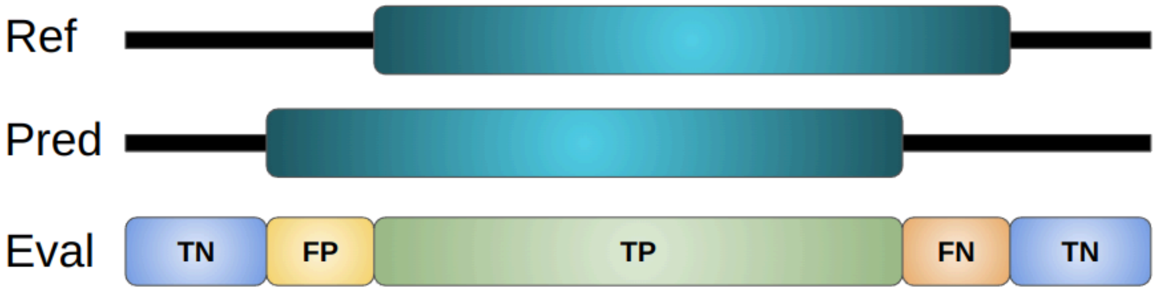


**Figure S2**. Schematic representation of the evaluative logic behind labeling the identified residues in the predictions compared to their associated reference.

| **Fold** | **Accuracy** | **Precision** | **Recall** | **F1-Score** |
|---|---|---|---|---|
| 1 | 0.88 | 0.91 | 0.91 | 0.9 |
| 2 | 0.89 | 0.92 | 0.93 | 0.91 |
| 3 | 0.89 | 0.93 | 0.91 | 0.91 |
| 4 | 0.87 | 0.9 | 0.91 | 0.89 |
| 5 | 0.86 | 0.9 | 0.91 | 0.89 |

**Table S5**. Performance metrics, calculated for each fold in the cross validation test, of the software in identifying STRP residues from non-STRP ones.

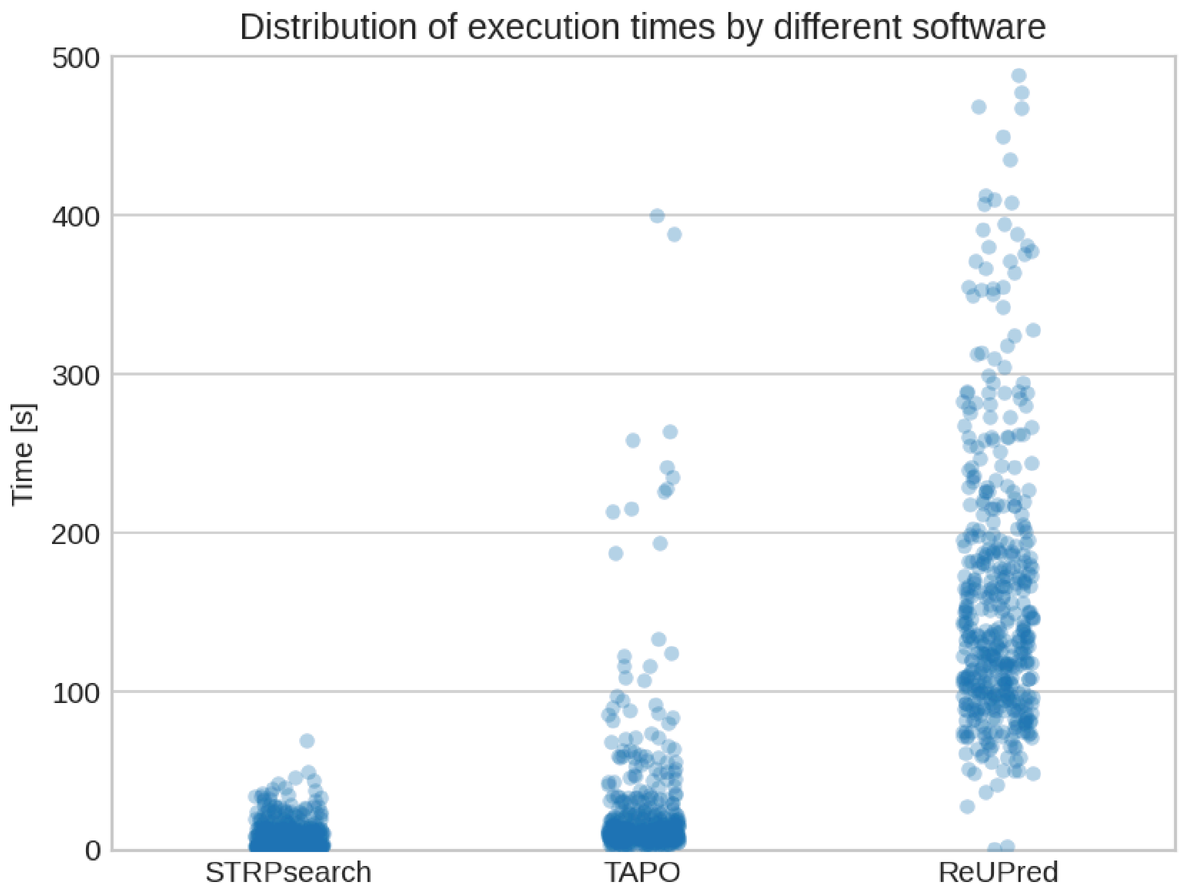


**Figure S3**. Distribution of execution times by different softwares on the benchmark dataset.
